# Pollination and abiotic stress alter the distribution of variation in floral longevity and opportunities for adaptation

**DOI:** 10.1101/2025.09.07.674481

**Authors:** Rachel B Spigler, Susanna Ostrowski

## Abstract

**Background and aims:** Floral longevity is thought to evolve by natural selection imposed by pollinators on heritable variation and resource constraints. In species with pollination-induced wilting, pollination rates should also influence phenotypic expression of genetic variation in longevity, raising questions of how and when variation in floral lifespan is shaped by adaptation.

**Methods:** We experimentally manipulated pollination rates in a wild population of *Sabatia angularis* (Gentianaceae). We asked how treatments influenced pollen gain curves, the phenotypic distribution of floral longevity, trait correlations, and potential costs. We further asked how abiotic factors influence longevity and how they may interact with pollination conditions.

**Key results:** Mean, variance, and maximum of longevity expressed increased with declining pollination. A flower longevity-number trade-off and positive longevity-flower size relationship persisted regardless of pollination context. Plants with more—not longer-lived—flowers suffered greater seed predation, potentially mitigating the flower number-longevity trade-off. The vapor pressure deficit (VPD) and seasonal changes constrained longevity. VPD had the strongest effect under the poorest pollination conditions, where longer-lived flowers would otherwise be favored.

**Conclusions:** Our findings highlight how ecological conditions and internal constraints shape phenotypic variation. Conditional expression of longevity may preserve genetic variation, enabling rapid evolutionary responses when environmental change causes selection and exposure of variation to align.

## 1 Introduction

It is well understood that environmental differences can present divergent selection pressures. However, for some traits, the environment governs not only the selective landscape but also the phenotypic expression of genetic diversity (Hoffmann & Hercus, 2000; Charmantier & Garant, 2005; Wilson *et al*., 2006; Ledón-Rettig *et al*., 2014). For example, stressful or extreme conditions might mask phenotypic variation, such as when a nutrient stress restricts plant growth despite underlying genetic variation in relative growth rates (Stanton *et al*., 2000). In other cases, variation may increase or only be exposed under extreme conditions, such as variation in water-use efficiency under drought (Edwards *et al*., 2012). Both scenarios can constrain adaptive evolution. However, in cases where variation is concealed under common conditions, genetic variation can be maintained, creating opportunities for adaptation when environmental change causes selection and exposure of variation to align.

Floral longevity—the length of time a flower remains open and functional—offers an opportunity to explore how ecological conditions influence selection and trait expression. Floral longevity varies dramatically across species, from less than a day to more than a month (Primack, 1985; Ashman, 2004; Song *et al*., 2022). Theoretical models used to explain this wide variation suggest that low or unpredictable pollination conditions should favor plants with longer-lived flowers because they maximize opportunities for pollen export and import, i.e., fitness gains (Primack, 1985; Ashman & Schoen, 1994; Schoen & Ashman, 1995; Charnov, 1996; Xu & Servedio, 2021). However, longer-lived flowers incur costs via resource demands of flower maintenance (Galen *et al*., 1993, 1999; Teixido & Valladares, 2014; Roddy *et al*., 2021) that are expected to result in a trade-off between flower number and longevity (Spigler & Woodard, 2019). Additionally, greater floral longevity may increase exposure to antagonists such as florivores, pathogens, or predispersal seed predators (Shykoff *et al*., 1996; Ashman, 2004; Teixido *et al*., 2011). As a result, shorter lived flowers should be favored when pollination conditions are reliable. Consistent with this framework, empirical studies have shown an inverse correlation between floral longevity and pollinator visitation rates across species or populations (Ashman & Schoen, 1996a; Arroyo *et al*., 2017). Yet pollination does not merely impose selection on floral longevity. In many species, it shortens floral lifespan by triggering petal wilting likely mediated through ethylene and other hormonal pathways (Stead, 1992; van Doorn, 1997). Consequently, observed (“realized”) floral longevity may not reflect intrinsic genetic potential longevity, and correlations between realized floral longevity and visitation rates may be driven, at least in part, by contemporary pollination conditions (Van Doorn & G, 1996; Lundemo & Ørjan Totland, 2007; Pacheco *et al*., 2016), rather than or alongside the outcome of selection. These considerations raise questions about how much variation in floral lifespan may be hidden within populations and under what conditions this variation is shaped by adaptation. They also highlight that much remains to be learned about the microevolutionary processes underlying the evolution of floral longevity.

Pollination-induced wilting could impact the evolutionary trajectory of potential floral longevity within populations by causing phenotypic variation, maintenance costs, and/or trait correlations to depend on the pollination environment. Studies have demonstrated how natural variation in pollination conditions can influence average flower lifespan via pollination-induced wilting in wild populations (e.g., Blionis & Vokou, 2001; Giblin, 2005; Blair & Wolfe, 2007; Castro *et al*., 2008), but it is unclear to what extent the distribution of phenotypic variation in longevity within a population is shifted vs. reshaped by those conditions. For example, the phenotypic distribution could be compressed under high pollination rates that push flowers to their minimum longevity, determined by the limits of plant signaling and cell death (Rogers, 2013). In this setting, realized floral longevity will be much shorter for genotypes with intrinsically long-lived flowers, but short-lived genotypes remain unaffected. Consequently, expression of additive genetic variation is reduced. Moreover, if there is genetic variation in wilting responses (the time from pollination to wilt), the rank order of genotypes might also change, affecting evolutionary outcomes. By changing the variance, the selective environment could alter trait correlations that influence the course of evolution (Sgrò & Hoffmann, 2004). Trade-offs might be diminished or obscured in high pollination environments, even when underlying intrinsic constraints remain, potentially lifting such limitations on genotypes with long-lived flowers. In contrast, phenotypic trade-offs with longevity could be strengthened or newly generated in environments where floral traits are under pollinator-mediated selection. Pollinator preference for more attractive plants (e.g., those with larger displays or larger flowers; Parachnowitsch & Kessler, 2010; Sletvold *et al*., 2010; Lavi & Sapir, 2015) could lead to faster pollination and earlier wilting, creating the appearance of a trade-off between attraction traits and floral lifespan. Because this trade-off is driven by pollinator preference and activity rather than or independently of intrinsic constraints, we term it a pollinator-mediated trade-off.

Abiotic factors can also effect plastic changes in floral longevity and alter the selective landscape independently of or jointly with the pollination environment. Indeed, others have highlighted how biotic and abiotic agents can interact to influence expression, selection, and evolution of floral traits more broadly (Caruso, 2006; Rusman *et al*., 2019; Day Briggs & Anderson, 2024). Multiple experimental studies have also shown that temperature and water or resource availability can influence longevity (Meagher & Delph, 2001; e.g., Vesprini & Pacini, 2005; Parra-Tabla *et al*., 2009; Arroyo *et al*., 2013; Jorgensen & Arathi, 2013; Spigler & Woodard, 2019). Resource availability can also affect the flower number-longevity trade-off (Spigler & Woodard, 2019). Hot and dry conditions, in particular, could influence the expression and fitness consequences of longevity by increasing floral transpiration rates and water loss (Teixido & Valladares, 2014; Bourbia *et al*., 2020). Paralleling the longevity-visitation correlation found across species, Song *et al*. (2022) demonstrated a negative correlation between longevity and temperature across species on a biogeographic scale, noting they could not distinguish the relative influences of genetics vs. plasticity. If abiotic stress stifles the expression of potential longevity, adaptive evolution of floral longevity could be constrained even if pollination conditions place strong selective pressure on longevity.

In this study, we experimentally manipulated pollination conditions to assess their influence the expression of floral longevity, phenotypic correlations, and potential costs of longevity in a wild population of the biennial *Sabatia angularis* (Gentianaceae) against a backdrop of natural abiotic variation. We ask the following sets of questions: (1) Do pollination conditions influence the distribution of floral longevity within a population, thereby shaping opportunities for adaptation? (2) Can we detect relationships between floral longevity and other floral traits in the field? Are relationships similar across pollination conditions, suggesting intrinsic correlations, or do they vary in a way that is more consistent with an ecological relationship mediated by pollinators (pollinator-mediated trade-off)? (3) How do abiotic factors influence the expression of floral longevity, and to what extent does this influence depend on pollination conditions? (4) Do plants with longer-lived flowers incur greater costs via predispersal seed predation, and, if so, are these costs modulated by the pollination environment? Beyond the evolution of floral longevity, the results of this study can provide broader insights into how internal and external constraints shape evolutionary potential and how variation within populations may be maintained.

## 2 Methods

### 2.1 Study species

*Sabatia angularis* is found in grasslands, roadsides, and outcrops throughout the mid-western and eastern United States. Plants flower from July to August, producing pink, nectarless flowers that are visited by a suite of generalist pollinators (mainly bees and flies). Flowers are protandrous, self-compatible, and contain ∼1000 ovules. Populations range from mixed mating to predominantly outcrossed, with outcrossing rate increasing with population size (Spigler *et al*., 2010). Seed production in populations can be pollen limited, with the degree of pollen limitation varying across years and populations (Dudash, 1993; Spigler, 2018). Plants can rely on autonomous selfing to reduce limitation to some degree under certain conditions (Spigler, 2018), but the process appears to be stochastic with no specific mechanism (Spigler & Maguiña, 2022). Pollinated flowers develop into dry dehiscent capsules from which seeds disperse passively. An unidentified, Lepidopteran predispersal seed predator is found in most populations to varying frequencies (Spigler, unpublished data). A single developing larva inside a developing fruit will eat all the seeds before exiting.

Prior work on floral longevity in *S. angularis* has found substantial intraspecific phenotypic variation in potential or “maximum” flower lifespan within populations, ranging from 3d to 21d, when grown under controlled, pollinator-free conditions (Spigler, 2017). Floral longevity is heritable and is plastic in response to resource availability (Spigler & Woodard, 2019). A trade-off between flower number and flower size and a positive correlation between lifespan and flower size were also found under controlled conditions, with evidence of a genetic basis for these relationships (Spigler & Woodard, 2019; Spigler & Charles, 2023). Pollen deposition, but not removal, induces wilting, causing realized longevity to be approximately half as long as maximum longevity (Spigler, 2017). Dudash (1991) noted flowers lasted only ∼4d in a wild population and that pollen removal can be rapid (∼90% pollen removed within 2d; Dudash, 1991). Flower number and size may be under pollinator-mediated selection (Emel *et al*., 2021), which could result in pollinator-mediated trade-offs with longevity.

### 2.2 Field study

This study was conducted in a large (>1,000 plants) population of *S. angularis* located in a serpentine grassland in southeastern Pennsylvania, USA (population ‘UB2’ Spigler, 2018). Prior to flowering, we randomly assigned plants to one of four pollination treatments (pollinated, control, emasculated, and bagged) to create a gradient of pollination rates within the same population. Pollinated plants received supplemental outcross pollen on all flowers using pollen available from other plants in the population. Control plants were not manipulated and experienced ambient pollination conditions. Emasculated plants had all anthers removed to reduce visitation rates to these plants (pollen is the sole pollinator reward in *S. angularis* flowers). Moreover, emasculation also eliminates the possibility of self-fertilization, making plants completely reliant on pollinators for seed production (no evidence of apomixis: Spigler, unpublished data). Emasculation does not damage *S. angularis* flowers; neither longevity nor seed set are reduced in emasculated flowers (Spigler, 2017, 2018). In addition, the type of pollen received (selfed vs. outcrossed) does not alter the effect of pollination on longevity (Spigler, 2017). Finally, bagged treatment plants were covered with cages made of bridal veil to exclude pollinators. Bagged plants were not emasculated for logistical reasons, namely because continued removal and replacement of bags can damage plants. In total, the study included 437 plants: 123 pollinated, 126 control, 119 emasculated, and 69 bagged.

We monitored plants three times per week to record the date the first flower opened. Once a plant began to flower, we tagged two flower buds for measuring floral longevity. We restricted tagging to flowers at secondary positions to control for potential position effects, though in some cases tertiary positions were used if all secondary flowers had already opened. We checked tagged flowers daily to record the date each opened and the date each wilted. We considered a flower to be wilted when all five petals had evidence of curling, as in Spigler & Charles (2023). We measured the length and width of a haphazardly chosen petal on at least 2 other flowers per plant and used the product of length and width as an estimate of flower size (as in Emel *et al*., 2021). At the end of flowering, we counted the total number of flowers produced, number of fruits, and number of fruits with evidence of seed predation (exit hole in fruit or larva still present inside). We collected predation data from 404 plants.

#### 2.2.1 Pollination rates

To confirm that the emasculation treatment was effective in reducing pollination rates, we conducted two studies. First, we observed pollinator visitation to control (N=20) and emasculated (N=31) plants. We used video cameras to record each plant for one hour between 9am and 3pm, with the order of filming randomized among plants. We subsequently quantified visitation using two metrics: visits per hour and number of visitors per hour. The former metric includes all visits per plant for a given visitor, whereas the second only counts the number of visitors. Visitors that left the field of view but may have returned were qualified as a new visitor. Visits were only counted when the insect contacted a flower.

Second, we quantified pollen deposition over time for control and emasculated plants. Because quantification requires destructive sampling (stigma removal), we identified an additional 48 plants to serve as control (N=24) and emasculated (N=24) plants. We tagged 7 buds per plant with jewelry tags and randomly assigned each a treatment corresponding to the age the stigma should be collected: 1, 2, 3, 4, 6, 8, and 10 days old. Tagged buds were checked daily. Once opened, we recorded the date and collected the stigma on the appropriate day using fine-tipped forceps, which were cleaned with 70% ethanol after each use. Stigmas were stored in 96-well plates at room temperature until processing. To quantify pollen loads, stigmas were soaked in 70% ethanol for 1-2 hours, mounted on a glass slide and viewed with a 4x objective lens on a fluorescence microscope with a green light filter. We captured photographs of the autofluorescent pollen and then counted the number of grains using the Cell Counter feature in ImageJ (Schneider *et al*., 2012).

#### 2.2.2 Abiotic factors

We obtained raw abiotic data collected from a meteorological tower located within the study population (described in Schedlbauer, 2015). These data included air temperature and relative humidity (CS215, Campbell Scientific, Logan, UT), precipitation (TE525, Texas Electronics, Dallas, TX), soil temperature (shielded Type-T thermocouples, Omega Engineering, Stamford, CT) and volumetric water content (“VWC”, EC-5, Decagon Devices, Pullman, WA) at 10 cm depth. Measurements were taken every 15 s and summed over 30 min intervals by a datalogger (CR 1000, Cambell Scientific). We used air temperature and relative humidity data from each interval to calculate vapor pressure deficit (VPD). VPD measures the deficit of moisture in the air relative to saturation and can drive transpiration rates. All else equal, high VPD conditions increase the rate of plant water loss, creating stronger physiological stress than low VPD conditions. Because soil temperature and VWC data were each collected from two sensors, we calculated the average measurement per interval across sensors. For each date, we determined cumulative precipitation and maxima for air temperature, relative humidity, VPD, soil temperature, and VWC. Finally, for each flower with recorded longevity measurements, we matched the dates it was open to corresponding abiotic data and calculated the average across the days representing the flower’s lifetime.

Ultimately, we focused on VPD as the potential main abiotic ecological driver of floral lifespan. Although VWC can be key in ameliorating water stress posed by high VPD, VWC varied minimally across the study period (0.238–0.260). In contrast, VPD ranged from 0.03 to 3.6 kPa, representing a strong gradient of atmospheric water demand. Together, these indicate that flowers were exposed to variable atmospheric stress across the season, but little drought stress. VPD was also highly correlated with temperature (air: r=0.84, p<0.0001; soil: r=0.80, p<0.0001).

### 2.3 Statistics

#### 2.3.1 Rates of pollinator visitation and pollen deposition

As proof of concept that emasculation served to create a reduced pollination rate condition, we first tested whether pollinator visitation was lower for emasculated plants compared with control plants. We adjusted visitation rate per plant to a per-flower basis by dividing each metric of visitation rate by the number of open flowers per plant on the day they were observed. Our two metrics, visitors per flower and total number of visits, were highly correlated (*r* = 0.87, *p* < 0.0001, N = 51). We elected to focus on visitor number because unusually high visitation by honeybees, which might inflate differences in visitation rate. We note that results are qualitatively the same for both measures (data not shown). This and all subsequent analyses were conducted using SAS^®^ OnDemand for Academics, Version 9.4, Copyright © SAS Institute Inc. For visualization purposes, figures were generated using R version 4.4.1 (R Core Team, 2024) within RStudio (Version 2024.09.1+394, Posit team, 2025) using the ggplot2 package (Wickham *et al*., 2016). We used a general linear mixed model to test whether visitors per hour varied as a function of pollination treatment and accounted for heterogeneity of variances between pollination treatments by including a random group term (proc mixed). We initially included a random term for day of observation, but removed it because it was not significant, affected model convergence, and reduced model fit.

We tested for differences in pollen deposition with flower age across pollination treatments using a general linear mixed model with repeated effects (proc mixed). We modeled the number of pollen grains per stigma (square-root transformed) as the dependent variable, with pollination treatment (emasculated vs. control), flower age, and their interaction as fixed variables. Patterns of pollen accumulation over flower lifetime were nonlinear, and each individual and treatment could be best described by different models. Therefore, we treated flower age as a categorical main effect, and we used prespecified contrasts to test for differences in pollen load between pollination treatments at each age. We included a random effect for plant identity. Although the order of sampling stigmas of different ages was randomized within each plant, pollen loads on two-day-old stigmas are more likely to resemble those on three-day-old stigmas than on ten-day-old stigmas. Therefore, we also used an autoregressive covariance structure to model pollen accumulation across time (age) within individual plants. Finally, we included the date the stigma was collected as a covariate to test and account for potential changes in pollen loads across the season.

For these and all following analyses, we tested model assumptions (normally distributed residuals, no significant heteroscedasticity among groups) and assessed presence of highly influential data points based on restricted likelihood distance, Cook’s Distance, covariance ratios, and/or the absolute value of studentized residuals. Although some points were identified as influential, their exclusion did not qualitatively impact the results; therefore, we included all data. We tested for significant heterogeneity of variances among pollination treatments with a likelihood ratio test comparing models that allowed treatment-specific variances to those assuming homogeneous variance. We retained the grouping structure when variance heterogeneity was significant.

#### 2.3.2 Impact of pollination environment, abiotic environment, and plant traits on floral lifespan

We tested the effect of pollination environment, represented by our pollination treatments, traits (flower number, flower size), and VPD on floral lifespan using a general linear mixed model (proc mixed). We also included the date each flower opened as a covariate to capture broad seasonal patterns. We z-scaled predictor variables to account for their different scales and facilitate comparison (Schielzeth, 2010) and added a quadratic term for VPD to address potentially non-linear trends observed in scatterplots. We included plant identity as a random subject term to account for repeated measures on the same individual.

We additionally included interactions between pollination treatment and all covariates to test specific hypotheses. The first set of hypotheses addresses whether any potential influence of flower number or flower size (estimated via petal area) on longevity is due to pollinator-mediated plasticity vs. underlying phenotypic (genotypic) correlations. If plants with more or larger flowers are more attractive to pollinators, they will be pollinated more quickly, triggering earlier wilting and causing a negative relationship between these traits and floral lifespan in treatments that allow natural variance in pollinator visitation (emasculated and control). Where pollinator influence is removed or standardized (bagged or hand-pollinated treatments, respectively), we do not expect this relationship. In contrast, if there is an intrinsic trade-off between flower number and floral longevity, there should be a negative impact of flower number on longevity across all treatments (although an interaction could arise if it less detectable or appears weaker due to truncation of variation in high pollination treatments). A genetic correlation between flower size and floral longevity should result in a positive relationship (based on Spigler & Woodard, 2019) in all treatments (though as for flower number, strength may vary). The second set tests whether abiotic conditions (VPD and/or date of flower opening) interact with the pollination environment to influence longevity. A significant interaction may suggest that stressful conditions could constrain potential floral longevity and thus adaptative evolution of longevity in pollinator-poor conditions.

In addition to testing for differences in mean among longevity across pollination treatments, we used complimentary approaches to evaluate differences in phenotypic distributions. First, we used the likelihood ratio test for significant heterogeneity of variances among pollination treatments in our general linear mixed model. Second, we conducted pairwise Kolmogorov–Smirnov (K-S) tests on mean-centered data (i.e., each observation minus its treatment mean) using R within RStudio. Removing differences in mean longevity allows the KS test to focus on differences in variance, skew, and kurtosis. We note that the K-S test is conservative when used with integer data (Noether, 1967), and we applied a Bonferroni correction to obtain adjusted p-values.

#### 2.3.3 Predispersal seed predation

We tested whether longer-lived flowers face a greater risk of predispersal seed predation. Given expected differences in floral longevity among pollination treatments, we predicted a significant main effect of pollination treatment on predation risk: highest in emasculated flowers, lower in control flowers, and lowest in pollinated flowers. Bagged flowers were included to serve as a control for predation: although they are expected to have the longest floral lifespan, bagging should exclude any predispersal seed predation occurring post-anthesis. We also anticipated an effect of individual-level floral longevity on predation risk. However, we expected a significant interaction between longevity and pollination treatment, potentially driven by the bagged treatment or by reduced variation in longevity within the more highly pollinated treatments. To evaluate these predictions, we used a hurdle model. First, we employed a generalized linear model to test whether the probability of predation (binary outcome: presence or absence of predation on any fruit) per plant was influenced by mean floral lifespan per plant, pollination treatment, and their interaction. We included flower number per plant as a covariate, as larger floral displays may attract more predispersal seed predators, and date of flowering, to capture any potential directional seasonal trends. Next, for plants that experienced at least one predation event, we used a binomial logistic regression to assess the effects of the same predictors on the proportion of fruits predated. By the time of fruit set, some plants had died, broken, had evidence of disease. We excluded these from our analysis for a total of 361 individuals, of which 108 (30%) had evidence of predation.

## 3 Results

### 3.1 Rates of pollinator visitation and pollen deposition

Pollinator visitation rate was significantly greater for control plants compared to emasculated ones (*F* = 9.37, *p* < 0.0001). Control plants had 1.8 times as many visitors per flower per hour (0.66±0.08SE vs 0.37±0.06SE). This difference translated into different pollen accumulation patterns, with the effect of pollination treatment on pollen deposition changing with flower age (Fig. 1, Table 1). No difference was observed on the first day flowers were open. However, contrasts indicated that pollen deposition on control plants quickly surpassed that on emasculated plants, maintaining higher pollen loads until the two converged on day 6, with no significant difference in pollen grains per stigma thereafter (Fig 1, Supporting Information Table S1). We also detected a negative effect of date on pollen deposition (Table 1), indicating average pollen loads declined across the season.

**Table 1.**
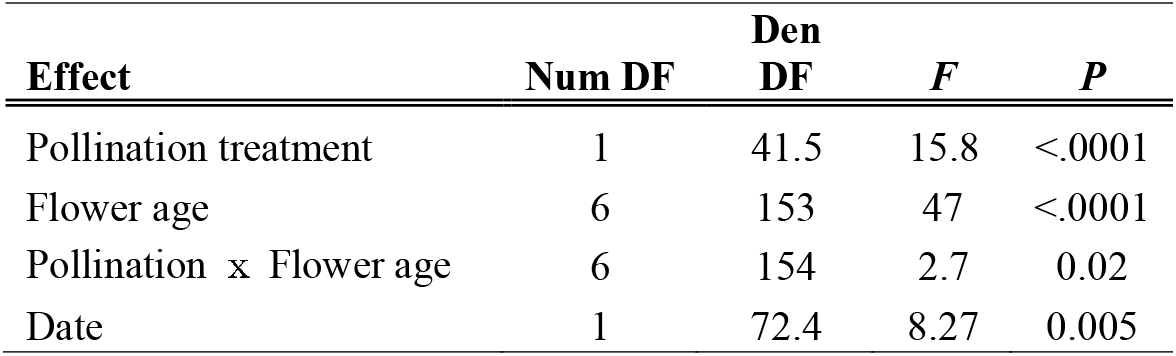
Results of mixed model testing the impact of pollination treatment, flower age, and their interaction on pollen deposition.

**Figure 1.**
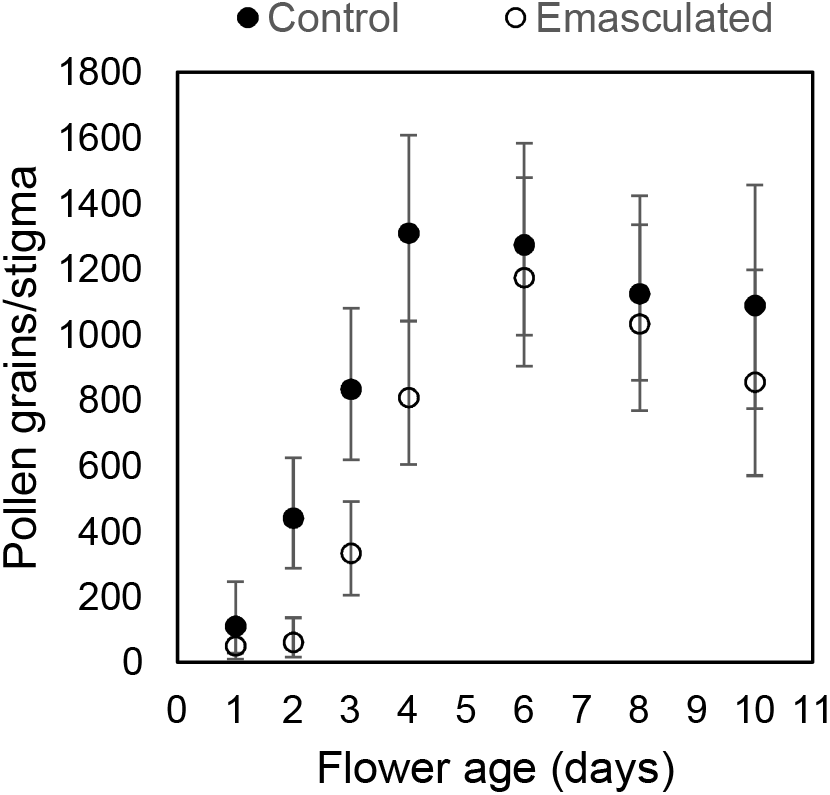
Pollen accumulation on stigmas with flower age. Marginal means and approximated 95% CI based on back-transformed data are plotted for control (solid black dot) and emasculation (open dot) treatments. Pollen deposition of emasculated plants is predicted to catch up to control plants when flowers are 6d old. See Tables 1 and SI1 for model results and pairwise comparisons between treatments per flower age sampled, respectively.

### 3.2 Impact of pollination environment, plant traits, and abiotic factors on floral lifespan

Pollination treatment influenced the mean, variance, and overall distribution of flower lifespan. Floral longevity increased with declining pollination rates: mean lifespan on pollinated, control, emasculated, and bagged plants were 4.1d (0.98SD), 4.5d (1.15SD), 5.0d (1.36SD), and 7.0d (2.2SD), respectively (Fig. 2) based on raw data. This effect was statistically significant (Table 2). In fact, all pairwise comparisons were statistically significant (Table S2). There was also significant heterogeneity of variances across pollination treatments estimated from the mixed model (*X*^2^= 65.9, df=3, p<0.0001, Table 3A), with variance similarly increasing with declining pollination conditions. This trend is weaker but also apparent from the coefficient of variation based on raw data: 0.24, 0.26, 0.27, 0.31 (pollinated<control<emasculated<bagged, respectively).

**Table 2.**
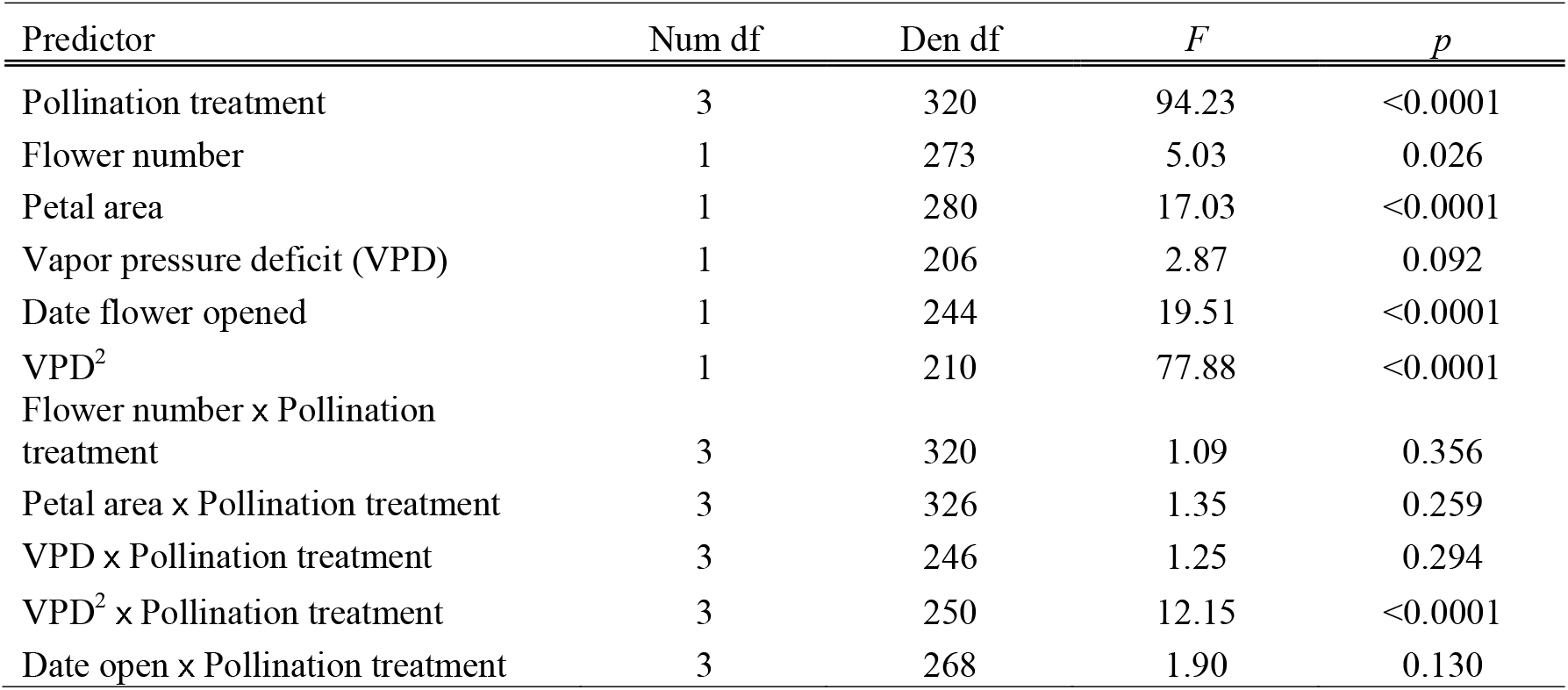
Results of mixed model testing the effects of pollination treatment, phenotypic covariates, and abiotic variation on mean floral longevity.

**Table 3.** Variance in floral longevity across treatments (A) and pair-wise distribution comparisons (B).

Heterogeneous residual variances ( ± SE)
| Treatment group | Estimate | SE |
| --- | --- | --- |
| B | 2.80 | 0.39 |
| E | 1.47 | 0.16 |
| C | 1.07 | 0.11 |
| P | 0.75 | 0.08 |

**B. Pairwise Kolmogorov–Smirnov test on centered values**
| Pairwise comparison | *D* | *p*<sub>adjusted</sub> |
| --- | --- | --- |
| B vs. E | 0.34 | <0.0001 |
| B vs. C | 0.27 | 0.0003 |
| B vs. P | 0.37 | <0.0001 |
| E vs. C | 0.18 | 0.0200 |
| E vs. P | 0.37 | <0.0001 |
| C vs. P | 0.25 | <0.0001 |

**Figure 2.**
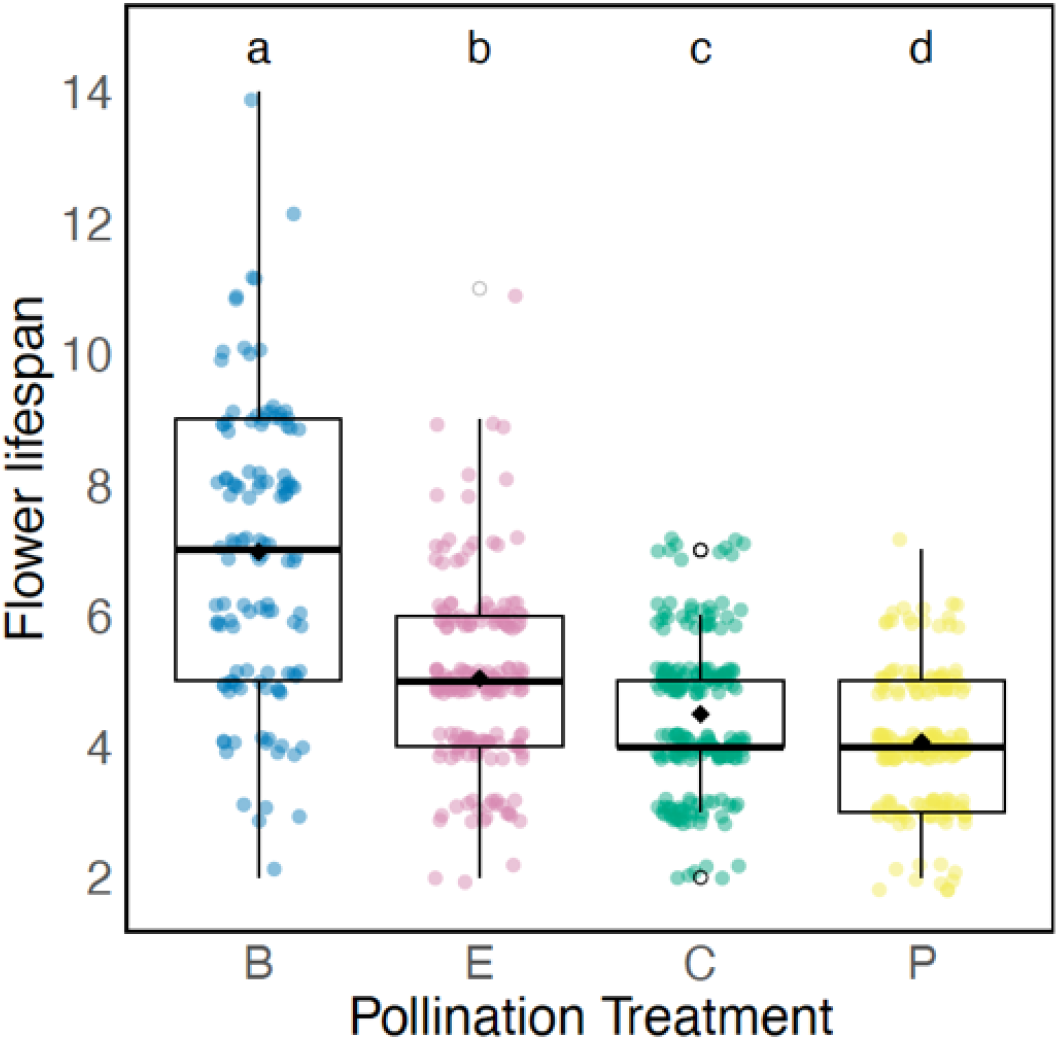
Box and whisker plots illustrating significant differences in mean, variance, and range of floral longevity across pollination treatments. Longevity is represented as lifespan of individual flowers in days. Treatments create a gradient of pollination rates declining from left to right. Treatment codes: B=bagged, E=emasculated, C=control, P=supplemental hand pollination. Raw data points were jittered to improve visibility. Different lowercase letters at top indicate significant differences in mean flower lifespan between treatments.

Differences in the phenotypic distributions are also evident in the range of phenotypes seen among treatments. Lifespans of individual flowers ranged from 2-7d in the pollination and control treatments. The lower limit was the same in the emasculated and bagged treatments, but the upper limit increased 1.57x in the emasculation treatment (range 2-11d) and doubled in the bagged treatment (range 2-14d). Pairwise comparisons of mean-centered distributions were also all significant (Table 3B), indicating that the shape and spread of the distribution of phenotypic variation in floral longevity depends on pollination treatments. These differences are also illustrated in the probability density and cumulative distribution functions (Fig. S1). The most dramatic difference can be seen in the distribution of longevity among bagged plants. The distribution is wider, and the weight of the distribution is to the left relative to the other treatments, with more extreme values above the mean (see cumulative distribution function, Fig. S1). This difference is less pronounced but visible and significant for the emasculation treatment.

Results from the mixed model further revealed that mean flower lifespan declined significantly with flower number (Fig. 3, Table 3). The interaction between pollination treatment and flower number was not significant, suggesting an intrinsic trade-off between flower longevity and number. Flower size, however, had a positive influence on longevity (Fig. 3, Table 3). This pattern is opposite to the predicted direction for a pollinator-mediated correlation and is instead consistent with a prediction of an intrinsic positive correlation. The effect appeared somewhat stronger in the bagged treatment, however the interaction between petal area and pollination treatment was not significant.

**Figure 3.**
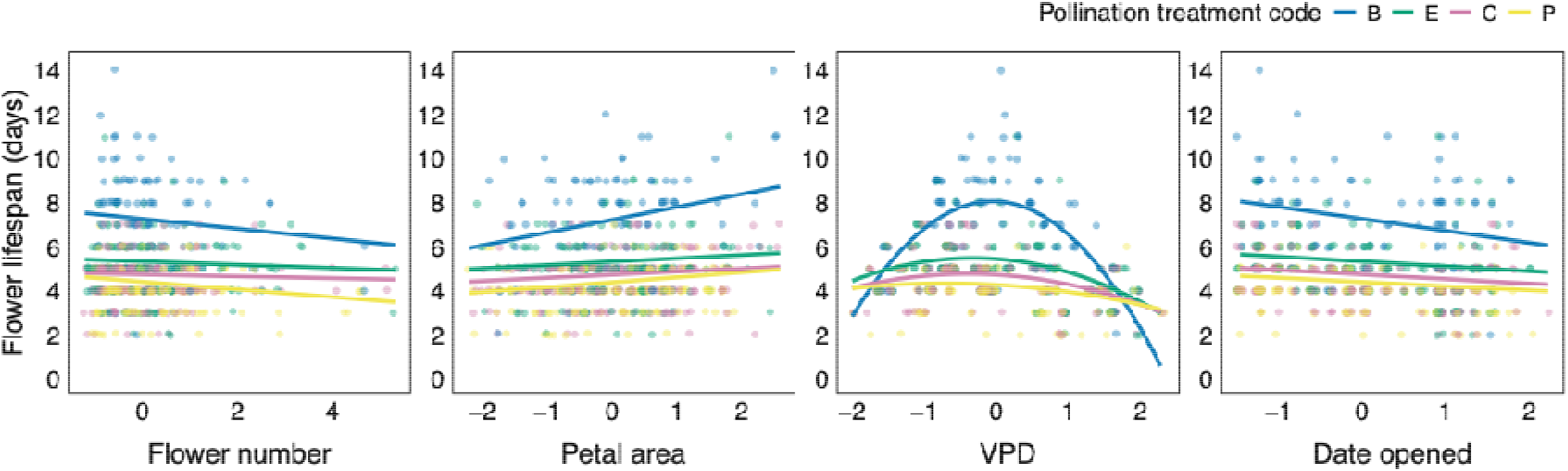
Effects of flower number, petal area, vapor pressure deficit (VPD) and date of opening on flower lifespan (days). Predictor variables were *z*-transformed (mean = 0, SD = 1; see Methods). Points show observed values, and lines represent model predicted relationships for each pollination treatment: B=bagged (blue), E=emasculated (green), C=control (pink), P=supplemental hand pollination (yellow). Points are jittered slightly in both axes for visibility.

The relationship between flower lifespan and VPD was non-monotonic. Flower lifespan was greatest around average VPD (Fig. 3C; significant quadratic term Table 2) and lowest at extreme conditions in both directions. The strength of the quadratic relationship, however, was statistically different across treatments (Table 2). The difference is most pronounced for the bagged plants; the estimated coefficient for the quadratic VPD term in the bagged treatment was between 3.6x and 10.6x greater than in the other treatments (Supplemental Table 2), illustrating greater potential for water stress to hamper adaptive evolution of longevity under poor pollination conditions. Viewed another way, all pairwise comparisons among pollination treatments are statistically significant at or near the mean VPD, but these differences diminish as VPD moves away from the mean in either direction. We also found that floral lifespan decreased across the season (Fig. 3, Table 3). Although the estimate of the slope is again greatest for the bagged treatment (between 2.9x to 7.9x that of the other treatments; Supplemental Table 2), the interaction between date opened and pollination treatment is not statistically significant (Table 3).

### 3.3 Seed Predation

Neither the probability of harboring a seed predator in at least one fruit nor the proportion of fruits predated (predation rate) depended on pollination treatment (Table 4). In fact, plants in the bagged treatment had similar predation rates to those found in the other treatments. Neither metric of seed predation was influenced by floral longevity, and the interaction between treatment and longevity was not significant (not shown). Date of flowering was not significant, indicating no directional seasonal patterns. However, both the presence of seed predation and its frequency significantly increased with flower number.

**Table 4.** Results of hurdle models testing the effects of pollination treatment flower lifespan, and first date of flower on predispersal seed predation.

| Predictor | Presence/absence |  |  |  | Predation rate |  |  |  |
| --- | --- | --- | --- | --- | --- | --- | --- | --- |
|  | Num df | Den df | *F* | *p* | Num df | Den df | *F* | *p* |
| Pollination treatment | 3 | 352 | 1.34 | 0.260 | 3 | 101 | 0.26 | 0.857 |
| Flower lifespan | 1 | 352 | 0.78 | 0.378 | 1 | 101 | 0.65 | 0.423 |
| Flower number | 1 | 352 | 7.21 | 0.008 | 1 | 101 | 19.34 | <0.0001 |
| Date of flowering | 1 | 352 | 0.56 | 0.455 | 1 | 101 | 0.81 | 0.369 |

## 4 Discussion

By simulating a gradient of pollination rates within a wild population and leveraging natural variation in abiotic conditions, we demonstrate how the ecological conditions that influence the selective landscape for floral longevity can also shape phenotypic expression of genetic variation. We also show that the vapor pressure deficit can substantially suppress the expression of maximum longevity, with the strongest impact in low pollination environments where longer floral lifespans could otherwise offer the greatest fitness benefit. In contrast, a trade-off between flower number and longevity and positive relationship between longevity and flower size were similar across pollination conditions, suggesting that certain costs and constraints may be intrinsic and persist regardless of the pollination environment.

### 4.1 Pollination conditions shape the phenotypic distribution of floral longevity and adaptive significance

Our experiment reveals that pollination rates influence mean realized floral longevity and the shape and spread of the distribution of phenotypic variation in *S. angularis*. In this way, our work aligns with previous studies demonstrating changes in mean longevity with pollination conditions as a result of pollination-induced wilting (e.g., Giblin, 2005; Blair & Wolfe, 2007; Castro *et al*., 2008; Arroyo *et al*., 2022) and advances our understanding by revealing how pollination dynamics can shape the evolutionary potential of longevity. In this way, our work fits within the context of understanding how the environment impacts heritability (e.g., Hoffmann & Merilä, 1999; Charmantier & Garant, 2005). We demonstrate that high pollination rates constrain the phenotypic expression of additive genetic variation in potential floral longevity in *S. angularis*. In fact, the top 50% of phenotypes expressed under suppressed pollination (emasculated) and pollinator exclusion (bagged) conditions are absent under control or supplemental pollination conditions. We suspect that opportunities for expression of even greater longevity exist under pollination conditions not simulated here. The maximum lifespan of 14d observed in our study was still lower than the 20d lifespan achieved by *S. angularis* flowers under controlled, pollinator-free conditions (Spigler, 2017; Spigler & Woodard, 2019). While our bagging treatment excluded pollinators, it allowed autonomous self-pollination, which also induces wilting once enough pollen falls onto the stigma (Spigler, 2017). This mechanism, albeit somewhat stochastic (Spigler & Maguiña, 2022), may provide reproductive assurance (Spigler, 2018). However, in some *S. angularis* populations, up to 90-100% of pollen within a flower is removed within the first two days (Dudash, 1991; Spigler & Ostrowski, unpublished data), making plants reliant on pollinators for pollen deposition. Emasculated plants in our study were entirely reliant on pollinators and received fewer visits than controls, but visitation rates were unusually high this year compared to others (Spigler, 2018), likely due to a placement of a new honeybee hive. Emasculated plants also represented a small proportion of a large population with abundant outcross pollen. Therefore, the opportunity for flowers to express full potential longevity may be greatest in small populations where pollen removal outstrips pollen deposition, provided resources are not limiting.

Direct estimates of phenotypic selection on potential longevity are difficult to obtain for species with pollination-induced wilting, as it creates something of a chicken-and-egg problem: plants observed to have short-lived flowers in the field could have high fitness either because shorter floral lifespans are inherently adaptive or because other floral traits increases pollinator attraction, leading to high pollen loads that, in turn, shorten lifespan. Nevertheless, our data on pollen accumulation rates demonstrate potential fitness consequences of short floral lifespans and how these consequences depend on the pollination environment. Flowers in the control treatment lasted on average ∼4.5d, corresponding to the age at which mean pollen loads surpass mean ovule number per flower. Under lower pollen deposition rates, represented by our emasculated treatment, only flowers that can last longer have the potential to achieve full seed set. Marques & Draper (2012) experimentally demonstrated this fitness effect by bagging flowers of *Narcisus sertinus* of different ages. They found that flowers that were bagged later had higher fruit set under pollen-limited conditions.

The opportunity for selection on longevity within *S. angularis* populations is likely variable; pollinator visitation, pollen deposition rates, and pollen limitation vary across years (Spigler, 2018). For example, pollen loads of only ∼500 grains/stigma were found in this population in 2014. In such years, living ∼0.5d longer as seen in our emasculation treatment is unlikely to be sufficient, and genotypes with short maximum floral lifespans could receive very few grains before wilting. Predictions about total relative fitness, however, are admittedly difficult. On the one hand, ovules can lose viability with age (Castro *et al*., 2008; Marshall *et al*., 2010; Marques & Draper, 2012; Spigler & Maguiña, 2022). On the other hand, greater longevity may instead serve to increase average fitness across all flowers per plant by increasing display size and pollinator attraction (Ishii & Sakai, 2001; Harder & Johnson, 2005; Teixido *et al*., 2019). Notably, display size (number of flowers open at a given time, on average) of *S. angularis* study plants in our study increased significantly alongside floral longevity across our simulated pollination gradient (data not shown). Even with the potential costs, waiting for pollinators may be the best or only option for reproductive in those populations where fast pollen removal or self-incompatibility limits or precludes reproductive assurance via autonomous selfing (Rathcke, 2003; Arroyo *et al*., 2006). The extreme levels of pollen limitation that enable selection and expression of the longest-lived flowers may be rare. However, if variation is hidden under weak selection in most years, that variation can be maintained, albeit subject to drift. In this way, our results on phenotypes parallel the concepts of cryptic genetic variation and conditional neutrality, where expression and fitness differences only occur in certain contexts (McGuigan & Sgrò, 2009; Anderson *et al*., 2013; Paaby & Rockman, 2014; Ledón-Rettig *et al*., 2014), contributing to the maintenance of variation and potentially rapid responses under intense selection.

### 4.2 Intrinsic and extrinsic limits on floral longevity across pollination contexts

#### 4.2.1 Trait correlations persist across pollination contexts

By changing the genetic variance, the selective environment can also influence trait correlations that are expected to affect the evolutionary trajectory of traits. We considered that this might apply to the pollination environment, either because the correlations are diminished in high pollination environments or because they are mediated by pollinators. The latter idea was also suggested by Brazeau *et al*. (2024). Even though pollination context affected the distribution of variation in longevity in our study, we did not detect significant differences in either the flower number-longevity trade-off or the positive relationship between longevity and flower size across pollination conditions. These patterns do not support the pollinator-mediated trade-off hypothesis and are instead consistent with intrinsic relationships. Prior work also supports a genetic basis for these relationships (Spigler & Woodard, 2019). Brazeau *et al*. (2024) similarly did not find evidence for a change in correlation between floral longevity and flower size across pollination treatments in *Mimulus guttatus*.

The trade-off between flower number and longevity is a key element in theory explaining the evolution of floral longevity as a resource allocation strategy (Ashman & Schoen, 1994). This marks the third time we have documented this trade-off in *S. angularis* (Spigler & Woodard, 2019; Spigler & Charles, 2023), and the first time it has been demonstrated in a wild population, providing compelling evidence that it is an evolutionarily relevant pattern. Identifying trade-offs within populations has proven elusive, but if found, can provide insight into the maintenance of genetic variation in fitness-related traits (Agrawal *et al*., 2010). A positive genetic correlation between flower size and longevity could also serve to maintain genotypes with high maximum longevity. When larger flowers are favored by pollinator-mediated selection (e.g., Lavi & Sapir, 2015) or fecundity selection (i.e., larger flower = more ovules; Mochizuki *et al*., 2019), an evolutionary response in flower size could lead to a correlated response in longevity.

#### 4.2.2 Abiotic stress reveals costs of longevity and limits the opportunity for selection under pollination limitation

In addition to pollination dynamics, water balance and physiological costs of longevity may play a crucial role in the evolution of floral longevity (Primack, 1985; Roddy *et al*., 2021). The significant effect of VPD on realized floral longevity shown in our study is consistent with the high costs of water stress found in other species (Galen *et al*., 1999; Lambrecht, 2013; Teixido & Valladares, 2014; Zhang *et al*., 2017) and suggest a substantial physiological toll of flower maintenance in *S. angularis*. High VPD places greater evaporative demand, with potentially substantial water loss via petals (Roddy *et al*., 2018; Carins-Murphy *et al*., 2023). In fact, water loss through flowers can be greater than that through leaves (Lambrecht, 2013; Roddy *et al*., 2018). Increased transpiration can lead to cavitation, turgor loss, and ultimately tissue damage or death (Bourbia *et al*., 2020; Carins-Murphy *et al*., 2023), strongly constraining the expression of maximum longevity. However, because temperature largely drives VPD and previous studies have linked shorter lived flowers with higher temperatures (Arroyo *et al*., 2013; Vega & Marques, 2015; Teixido & Valladares, 2015), it is also possible that high temperatures directly reduced the longevity of *S. angularis* flowers in our study.

Unexpectedly, we found that longevity was also suppressed when VPD was much lower than average. Evidence of nonlinearity was strongest in the bagged treatment, but still visible and significant in all other treatments. Low VPD is typically associated with cool and humid conditions, but raw values of mean maximum VPD across floral lifespan in our study ranged from moderate to high (0.9kPa to 2.0kPa). Flowering in *S. angularis* occurs in late July and throughout August, when temperatures are consistently high, but rain is inconsistent. It is possible that high humidity in the lower range of VPD seen in our study could have reduced evaporative cooling and increased heat load on floral tissues. This effect was experimentally demonstrated Ipomoea species, wherein the temperature of corollas and gynoecium were significantly higher on flowers that were prevented from transpiring (Patiño & Grace, 2002). Other factors could have contributed to the unexpected quadratic relationship. For example, VPD determines the atmospheric demand for water, but soil water availability determines how quickly plants can replace lost water and can directly impact floral lifespan (Caruso, 2006; Jorgensen & Arathi, 2013). Yet, volumetric soil water content did not vary substantially across the season. Water stress created by VPD could also influence plant-pollinator interactions (Descamps *et al*., 2021; Kuppler & Kotowska, 2021) that in turn could alter patterns of pollination-induced wilting. For example, greater pollinator visitation at low VPD could underlie shorter floral lifespans under those conditions. We also note that metrics other than maximum daily VPD, such as daily integrated VPD, could serve as a better predictor but also may lead to a similar pattern (e.g., Roddy & Dawson, 2013). Regardless, our study shows a clear impact of VPD on floral longevity.

Expression of longevity in *S. angularis* is also increasingly constrained as the flowering season advances. The longest-lived flowers (up to 14d) in the study were only found at the start of the season, and mean longevity decreases by approximately 25% from the start to end of the season, based on our model. Other studies have noted seasonal effects on longevity within populations, though not necessarily in the same direction (Prokop *et al*., 2021; Lee & Caruso, 2022). Prokop *et al*. (2021) suggested that greater longevity of common chicory (*Cichorium intybus*) flowers later in the season may be driven by declines in temperature and day length from August to September. Temperature would not explain the decreased longevity across the season in our study, as there is no directional temperature change during this time. While we cannot identify the specific driver of the seasonal pattern in our study, changes in abiotic conditions are more likely at play than changes in biotic interactions. Although we found a seasonal trend for pollen deposition, pollen deposition decreased across the season, which in turn would lead to longer, not shorter, flower lifespans on plants that flowered later. In addition, there was no seasonal trend in predispersal seed predation, and herbivory is not common in this population (personal observation).

Both variables, but most substantially VPD, constrained longevity such that differences in mean longevity across treatments were eliminated or decreased. The implication is that this could prevent adaptive evolution under the poorest pollination conditions, where we expect selection to favor longer-lived flowers. These results mirror those found by Caruso (2006), wherein drought limited expression of some traits that would otherwise have been favored by pollinator-mediated selection. There could still be an evolutionary pathway to greater longevity under water stress if genetic variation in transpiration efficiency exists in the population (Vadez *et al*., 2014). It is also important to note that reduced longevity in response to temperature and water stress can save plants from future costs that would otherwise be put into flower maintenance, ultimately resulting in greater seed production in some pollen-limited populations (Dudley *et al*., 2018).

#### 4.2.3 Greater floral longevity does not increase risk of predispersal seed predation

Seed predation has the potential to constrain the evolution of greater longevity, but to our knowledge has been examined in only one other study (Fenner *et al*., 2002). We found that the cost of seed predation was paid by plants with more flowers, not longer-lived ones. Predation rates were also the same across our pollination treatments, including bagged plants. This finding suggests that oviposition occurs at the flower bud stage. While we don’t yet know the exact species, we know that our predator is a type of moth, and some Lepidopteran species oviposit in or on flower buds (Duggan, 1985; Thompson & Pellmyr, 1991). Fenner *et al*. (2002) demonstrated this type of predation in many Asteraceae species and, likewise, did not find an association between predation and capitulum longevity. Our finding of a strong effect of flower number on predation risk has also been reported in other studies (reviewed in Kolb *et al*., 2007). Given the potential for seed predation to reduce fitness, its association with increased flower number in *S. angularis* might even mitigate the trade-off with longevity observed in all pollination environments.

### 4.3 Conclusions

Our study highlights that understanding where and how adaptive evolution of floral longevity occurs requires disentangling the effects of extrinsic and intrinsic factors on the distribution of variation in floral lifespan. The impacts of pollination environment and abiotic factors on floral longevity shown here may become even more relevant as global change continues to alter pollinator availability, temperature, and water availability (Day Briggs & Anderson, 2024). Direct estimation of selection gradients on longevity may be challenging in species that exhibit pollination-induced wilting, but our findings illustrate how abbreviated flower lifespans could limit female fertility. Selection on maximum longevity in these species might occur infrequently, depending on the level of pollen limitation. However, if long-lived phenotypes are typically suppressed, this may allow genetic variation to be maintained, enabling episodic periods of selection in years of severe pollination that could drive rapid evolutionary change. Moreover, the induced wilting response may also be under selection (Ashman & Schoen, 1996b; van Doorn, 1997; Trunschke & Stocklin, 2017). Ultimately, this work underscores the need to better understand the environmental conditions under which floral longevity is expressed, constrained, and selected to predict how often and under what circumstances it can evolve via selection.

## 5 Acknowledgements

The authors thank Natural Lands for permission to work on their property. Estimating longevity requires checking plants 7 days a week over 6 weeks or more. With hundreds of plants in the field, this work would not have been possible without Matthias Gaffney and an amazing team of undergraduate researchers: A. Woodard, M. Baltazar, D. Bloom, J. Capista, M. Landy, I. Ortiz, and J. Wible. We also thank Jessica Schedlbauer, who generously provided us with raw abiotic data. This work and participant support was funded in part by Temple University and grant NSF DEB-1655772 awarded to RBS.

## 6 Author contributions

RBS conceived and designed the study, and was responsible for data collection, analysis, and interpretation. SO led the pollen study, including protocol development, data collection, and interpretation, and assisted with data collection for the rest of the study. RBS prepared the initial manuscript draft, with edits by SO. Both authors approved the final version of the manuscript.

## 8 Supporting Information

**Figure S1** Smoothed probability density and cumulative distribution function plots of observed floral longevities under four pollination conditions.

**Table S1** Pairwise comparisons of pollen accumulation between pollination treatments per flower age.

**Table S2** Pairwise comparisons of mean floral longevity between pollination treatments

## Notes

### Competing Interest Statement

The authors have declared no competing interest.

