## Supporting Information for "Pollination and abiotic stress alter the distribution of variation in floral longevity and opportunities for adaptation"

The following Supporting Information is available for this article:

**Fig. S1** Smoothed probability density and cumulative distribution function plots of observed floral longevities under four pollination conditions.

**Table S1** Pairwise comparisons of pollen accumulation between pollination treatments per flower age.

**Table S2** Pairwise comparisons of mean floral longevity between pollination treatments.

**Fig. S1** Smoothed probability density and cumulative distribution function plots of observed floral longevities under four pollination conditions. The distribution of variation in floral longevity varies significantly across pollination treatments (B=bagged, E=emasculated, C=control, P=pollinated). The most dramatic difference can be seen in the distribution of longevity among bagged plants: the distribution is wider, and the weight of the distribution is to the left with more extreme values above the mean (see CDF) relative to the other treatments. This difference is less pronounced but visible and significant for the emasculation treatment.

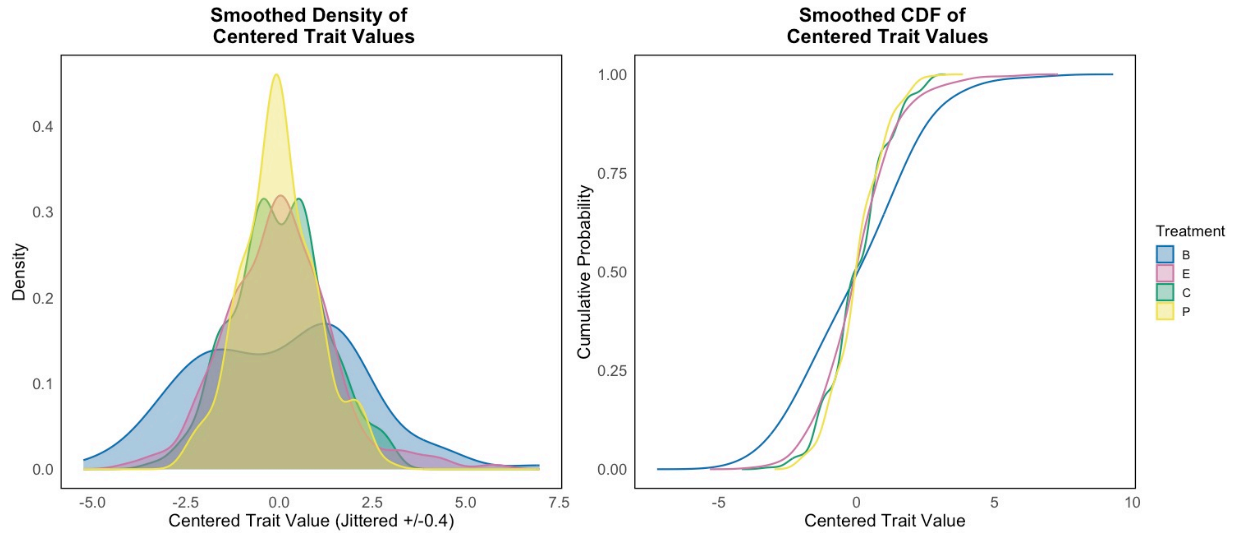


**Table S1** Pairwise comparisons of pollen accumulation between pollination treatments per flower age.

| Flower age (days) | Num DF | Den DF | *F* | $p_{adjusted}$ |
|---|---|---|---|---|
| **1** | 1 | 259 | 1.09 | 0.30 |
| **2** | 1 | 243 | 21.72 | <.0001 |
| **3** | 1 | 244 | 13.92 | 0.0002 |
| **4** | 1 | 241 | 7.76 | 0.006 |
| **6** | 1 | 248 | 0.23 | 0.63 |
| **8** | 1 | 250 | 0.21 | 0.65 |
| **10** | 1 | 258 | 0.99 | 0.32 |

**Table S2** Pairwise comparisons of mean floral longevity between pollination treatments.

| Treatment code | Treatment code | Estimate | SE | DF | *t* | $p_{adjusted}$ |
|---|---|---|---|---|---|---|
| **B** | **E** | 1.6308 | 0.1915 | 169 | 8.52 | <.0001 |
| **B** | **C** | 2.2263 | 0.1855 | 152 | 12 | <.0001 |
| **B** | **P** | 2.5843 | 0.181 | 139 | 14.28 | <.0001 |
| **E** | **C** | 0.5954 | 0.1174 | 353 | 5.07 | <.0001 |
| **E** | **P** | 0.9535 | 0.1102 | 324 | 8.65 | <.0001 |
| **C** | **P** | 0.3581 | 0.09918 | 363 | 3.61 | 0.002 |
